# Huntingtin CAG repeat size variations outside the Huntington’s disease complex: associations with depression and anxiety phenotypes and basal ganglia structure

**DOI:** 10.1101/2024.05.22.595390

**Authors:** Magdalena Vater, Nicolas Rost, Gertrud Eckstein, Susann Sauer, Alina Tontsch, Angelika Erhardt, Susanne Lucae, Tanja Brückl, Thomas Klopstock, Philipp G. Sämann, Elisabeth B. Binder

**Author notes:** Equal contribution (Philipp G. Sämann, Elisabeth B. Binder). **Corresponding authors information:** Magdalena Vater, MD; Elisabeth Binder, MD PhD, Prof.; Philipp G. Sämann, MD, E-Mail.

## Abstract

Huntington’s Disease (HD) is strongly associated with psychiatric symptoms, yet, associations between Huntingtin gene (*HTT*) CAG repeat size variations and psychiatric phenotypes outside the HD complex are still under-investigated. In this genetic case-control study we compared the distribution of *HTT* CAG repeat sizes in predefined ranges between patients with major depressive disorder (MDD) (n=2136) and anxiety disorders (ANX) (n=493), and healthy controls (CON) (n=1566). We used regression models to study interactions between the alleles and associations with fine-granular clinical phenotypes and basal ganglia structure. HD mutations in the range of incomplete penetrance (36-39 repeats) were not overrepresented in patients. In participants older than 48 years, 13-20 repeats on both *HTT* alleles were associated with a reduced ANX risk whereas a 13-20|21-26 combination was associated with an increased ANX risk. Post-hoc analyses confirmed a turning point around 21 repeats and trends in the same direction were detected for MDD. The joint patient|CON analysis of the full spectrum of allele combinations confirmed interaction effects and age-dependent allele|risk profiles. A short-by-long interaction effect and an age-dependent negative correlation of the short allele on the nucleus accumbens volume was detected, independently of the diagnostic group. In conclusion, we revealed that *HTT* CAG repeat sizes of both alleles in the non-HD range modulate the susceptibility for common psychiatric disorders and basal ganglia structure in an age-dependent way, displaying that normal variation of the functionally diverse wildtype huntingtin protein may already impact brain function.

## INTRODUCTION

The *HTT* gene represents the key locus responsible for the pathogenetic cascade occurring in Huntington’s disease (HD). It lies on Chromosome 4p16.3, and a pathologic elongation of the CAG repeat stretch in its first exon causes the dominant genetic disorder HD^1–3^. Clinically, HD leads to a progressive hyperkinetic movement disorder, cognitive decline and other behavioural abnormalities^1,4^ such as commonly reported apathy but also mood disturbances, anxiety, disinhibition, perseveration, psychotic symptoms and suicidality^5–8^. Still, the relevance of *HTT* CAG count variations for psychiatric symptoms outside the HD disease complex remains largely unclear. Currently, genome-wide association studies (GWAS) represent the prevailing tool to approach the genetics of psychiatric disorders, yet, they cannot directly measure variably expanded DNA repeats to clarify the significance of specific *HTT* CAG repeat sizes for psychiatric risk^9–11^.

The risk to develop HD is dependent on the repeat size. While alleles up to 35 *HTT* CAG repeats are viewed as not contributing to individual disease risk, there is a risk of anticipation with alleles in the range of 27–35 that are associated with a possible further elongation of the *HTT* CAG stretch in the next generation, especially with paternal transmission. Repeat sizes between 36 and 39 *HTT* CAGs can be found in individuals both affected or unaffected by HD, indicating incomplete penetrance. Alleles with 40 or more repeats will inevitably cause HD within the normal life span^12,13^. The age at onset of motor symptoms usually lies in mid-life and is negatively correlated with the length of the *HTT* CAG stretch on the HD-allele and further moderated by other genetic and lifestyle factors^14^.

The pathological hallmark of HD is a degeneration of the striatum that – when quantified by image segmentation – represents a sensitive biomarker for the disease^15,16^. Morphological differences of the putamen and caudate occur early, already in non-symptomatic, premanifest stages of patients with the pathogenic mutation^16–18^.

Overall MRI volumetry is considered more sensitive than clinical scoring^18,19^ to classify disease progression. The nucleus accumbens and pallidum are affected in the premanifest stage^16^, and later atrophy spreads to the insular and other cortices^20^. Basal ganglia volumes are predicted by the excess of the *HTT* CAG repeat expansion in conjunction with age but also corticostriatal pathways degenerate in correlation with the genetic load^21,22^. One study found that HD patients with an onset with psychiatric symptoms (as compared to patients with motor symptoms at onset) were younger, showed higher CAG repeat numbers and more neuronal density changes in the nucleus accumbens^23^.

In Western populations, the prevalence of HD estimates ranges from 10.6-13.7/100000^1,24–26^. As recently suggested^12^, the overall prevalence of *HTT* CAG expansions in the HD range could be even higher, with a prevalence of HD carriers of approximately 1/400, mainly carried by expansions in the HD range but of reduced penetrance. Perlis et al.^27^ reported *HTT* CAG expansions in the lower range of potential HD disease risk to be overrepresented in major depressive disorder (MDD), suggesting that psychiatric symptoms etiologically attributable to HD may ‘mimic’ symptoms of MDD or that an otherwise increased risk of MDD might be associated with this CAG repeat range. Another study revealed two cases with expanded alleles (both 37 *HTT* CAG repeats) among 2165 subjects of two cohorts with depressive disorders compared to no cases among 1058 control subjects^11^. The same study also claimed a nonlinear risk modulation of lifetime depression by *HTT* CAG repeat lengths, even in the non-HD range. The possibility that *HTT* CAG repeat lengths in the non-HD range potentially play a role in psychiatric disorders is further corroborated by reports on two large observational HD cohorts that pointed out an association of the presence of psychiatric symptoms with repeats in the range of 27–35 compared to lower counts^28,29^.

Here we investigated the distribution of *HTT* CAG repeat lengths in MDD and ANX compared with healthy subjects in order to calculate contributions to the disease risk and to study effects of *HTT* CAG repeats on selected dimensional psychiatric phenotypes and basal ganglia structure. A special focus was laid on *HTT* CAG repeat lengths of both alleles, their interaction and the impact of age.

## METHODS

### Source samples and final sample composition

We combined data from the Recurrent Unipolar Depression (RUD) study, a cross sectional case-control study on patients with recurrent unipolar depression and control subjects^30–32^ and the Munich Antidepressant Response Signature (MARS) project with subprojects *MARS-Depression*, a prospective multi-center naturalistic observational study of treatment outcomes in acutely depressed in-patients^33^, *MARS-Anxiety*, a study consecutively recruiting from the Anxiety Disorders Outpatient Clinic at the Max Planck Institute of Psychiatry (MPIP) ^34,35^, and *MARS-Controls*, a population cohort study randomly selected from the Munich resident’s registry^36^. Detailed sample descriptions, inclusion and exclusion criteria and respective diagnostic instruments are detailed in the **Supplementary Data**.

Overall, 4212 subjects from the RUD-study and MARS project were eligible for genetic analyses. For 10 subjects (6 MARS-Depression, 2 MARS-Anxiety, 2 RUD controls) no DNA sample was available and in 7 subjects (6 RUD cases, 1 RUD control) *HTT* genotyping failed due to poor sample quality, leaving 4195 subjects for further analysis of whom 2136 had MDD (58.5% women, age: mean [SD] 49.2 [14.1] years, range 18-87 years; ICD-10: F32: 20.2 %, F33: 79.8%), 493 had ANX as main diagnosis (58.2% women, age: mean [SD] 37.7 [12.1] years, range 17-75 years; ICD-10 diagnoses: F40.0: 1.8%, F40.01: 60.0%, F40.1: 9.3%, F40.2: 2.6%, F40.9: 0.2%, F41.0: 21.9 %, F41.1: 2.8% and F41.2: 1.2%) and 1566 individuals were control subjects (63.0% women, age: mean [SD] 49.6 [13.8] years, range 18-90 years).

### Molecular methods

*HTT* CAG repeat sizes were quantified by applying a modification of the method reported by Batsepe and Xin^37^. A PCR was performed using the primers CAG1-Met-Fwd 6-Fam-ATGAAGGCCTTCGAGTCCCTCAAGTCCTTC and CAG2-Rev GGCGGTGGCGGCTGTTGCTGCTGCTGCTGC (Metabion). The fragments were analyzed on an ABI 3730 DNA Analyzer (Applied Biosystems) with GeneScan500 ROX (Applied Biosystems) as internal size standard and results were processed by GeneMapper™ Software 5 (Applied Biosystems). The fragment sizes were converted to *HTT* CAG repeat numbers by comparison to predetermined *HTT* CAG repeat numbers of standard DNA (Standard Reference Material® 2393, National Institute of Standards & Technology, USA). Pathologic results (*HTT* CAG repeats >35) were confirmed by replication. **Supplementary Fig. 1** shows examples of the fragment analysis. More technical details are given in the **Supplementary Data**.

**Figure 1.**
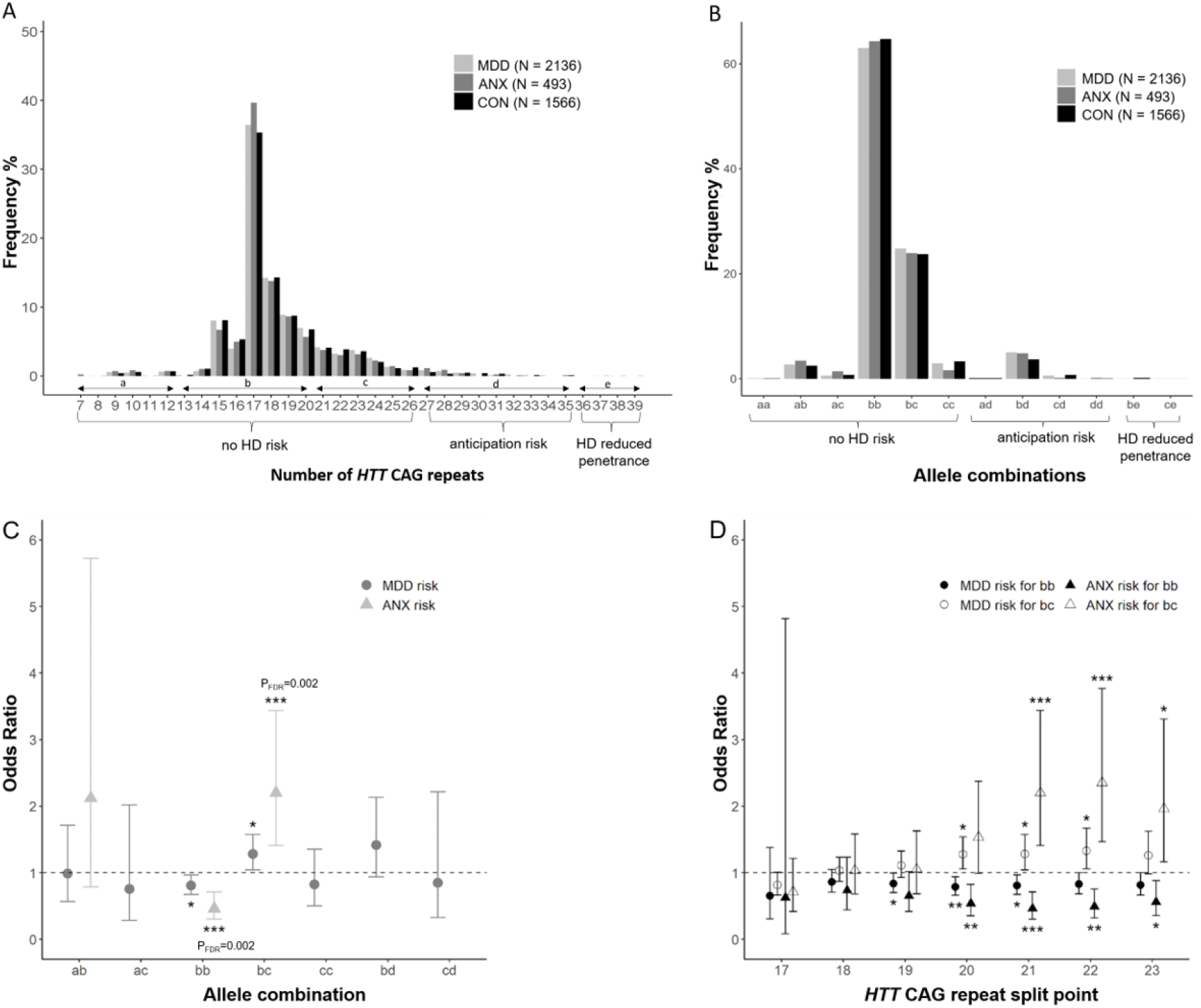
Distribution of *HTT* CAG repeat counts and association of MDD and ANX risk. Frequency distribution of the *HTT* CAG repeat counts in MDD, ANX and CON. **(A)** Bins represent *HTT* CAG repeat counts, considering both A1 and A2. Note canonical allele ranges are given as levels of the x-axis: *d* (27-35 repeats) and *e* (36-39 repeats), in addition to the ranges defined for this study: *a* (7-12 repeats), *b* (13-20 repeats) and *c* (21-26 repeats). **(B)** Bins represent all available combinations of the A1 and A2, each classified into ranges *a*–*e*. **(C)** ORs were calculated for MDD and ANX versus CON for different allele combinations in subjects ≥ 48 years. Error bars indicate 95% confidence intervals. ORs were not calculated for allele combination with a frequency of less than 5. *, ** and *** indicate nominal P-values < 0.05, < 0.01 and < 0.001 respectively. Only FDR corrected p-values < 0.05 are indicated as numbers. For comparisons between MDD (N for *ab* 30; *ac* 8; *bb* 739; *bc* 302; *cc* 34; *bd* 68; *cd* 9) and CON (N for *ab*: 23; *ac*: 8; *bb*: 604; *bc*: 189; *cc*: 31; *bd*: 37; *cd*: 8) ORs of allele combinations *ab, ac, bb, bc, cc, bd* and *cd* are shown. For comparisons between ANX (N for *ab*: 5; *bb*: 46; *bc*: 35) and CON (N= see above) ORs of allele combinations *ab, bb* and *bc* are shown. **(D)** ORs for allele combinations *bb* and *bc* for MDD vs. CON and ANX vs. CON in subjects ≥ 48 years over different category ranges are shown. The x-axis indicates variable split points between categories *b* and *c*. For instance, given a split point of 21, category *b* includes CAG repeat sizes of 13-20 and category *c* of 21-26; *, ** and *** indicate nominal P-values < 0.05, < 0.01 and < 0.001 respectively. Only FDR corrected p-values < 0.05 are indicated as numbers.

### Magnetic resonance imaging (MRI) samples and processing

The structural MRI sample composition and processing as well as the inclusion of an HD sample are detailed in the **Supplementary Data**.

### Statistical analysis

*HTT* CAG repeat counts on both alleles of one subject are referred to as A1 (lower count) and A2 (equal or higher count). When multiple tests of the same hypothesis were applied, the false discovery rate (FDR) method (q<0.05) was used^38^. All statistical analyses were performed in R version 4.0.3^39^, and all figures were created in R or MATLAB version v9.2.0, R2017a.

#### Comparing mean HTT CAG repeat counts and repeat size categories

Direct correlations between age or sex and the repeat counts were explored in MDD, ANX and CON (**Supplementary Tab. 1**). Differences of mean *HTT* CAG repeat counts between CON and patient groups were analyzed by separate one-way analyses of variance (ANOVA) (**Supplementary Tab. 2**) and by one-way analyses of covariance (ANCOVA) with age and sex as covariates (**Supplementary Tab. 3**). To compare frequency distributions between patients and control subjects for predefined combinations of *HTT* CAG repeat ranges *a* (7-12), *b* (13-20), *c* (21-26), *d* (27-35), *e* (36-39) – with resulting twelve combinations (*aa, ab, ad, ad, bb, bc, bd, be, cc, cd, ce, dd)* – we applied Fisher’s exact test **(Supplementary Tab. 5)**. Further, for those combinations with a minimum of five subjects in the respective cells, odds ratios (ORs) were calculated both for the whole sample and the age-split subsamples (**Fig. 1, Supplementary Fig. 5, Supplementary Tab. 6**). Post-hoc, given risk effects in the categories *bb* and *bc*, age and sex differences between the respective patient and CON groups within the combination *bb* and *bc* were analyzed (**Supplementary Tab. 4**.**1)**. In addition, age and sex differences *between* these combinations were analyzed in CON to exclude that risk effects were based on the age or sex stratification of the patient samples (**Supplementary Tab. 4**.**2**).

#### Combined linear and nonlinear influence of allele A1 and A2 and their interaction

To explore the combined influence of both alleles on psychiatric risk, pooling MDD and ANX, we estimated binomial logistic regression models with five terms A1, A2, A1^2^, A2^2^ and A1-by-A2 as predictors for the younger and older subjects (N=4195, median split point 48 years). Allele counts A1 and A2 were correlated (*r*=0.36; *p*<0.001; **Supplementary Fig. 3A**), so we accounted for collinearity by performing Gram-Schmidt-orthonormalization of all five terms. To investigate age interaction effects, a common model with all subjects and a two-level factor *age* was estimated. Relative disease risks were calculated by dividing the risk probability by the original patient|CON ratio for every allele combination represented in the sample. Resulting values > 1 thus represent an increased risk for MDD or ANX. Color-coded 3D-planes were plotted to visualize the relative risk for all A1|A2 combinations (**Fig. 2**), and 3D-histograms were added to illustrate the actual data density of the A1|A2 combinations (**Supplementary Fig. 2**).

**Figure 2.**
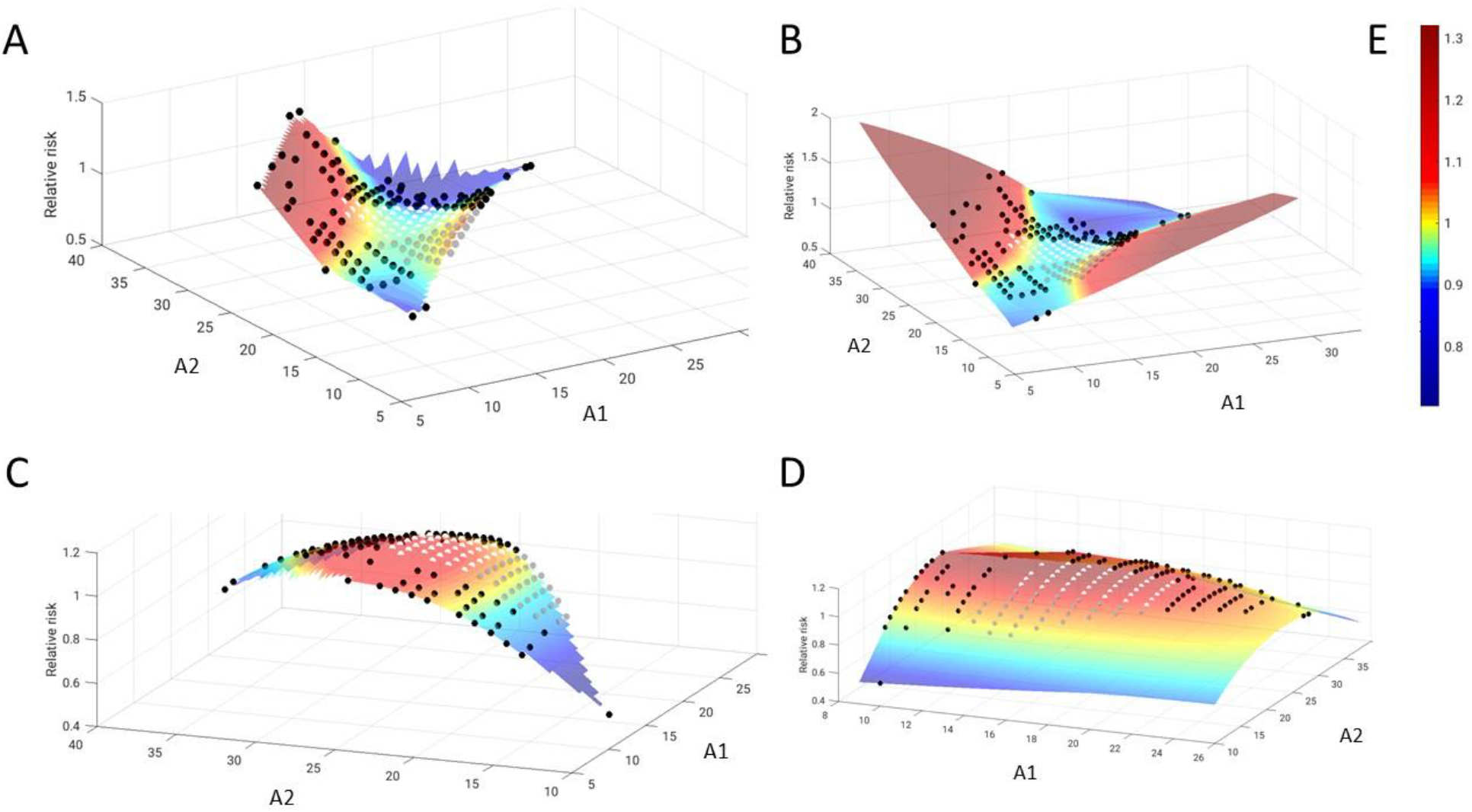
Graphical depiction of the relative psychiatric risk depending on both alleles and their interaction. For the full underlying statistical model see methods section. To allow for a comparison with the categorized repeat lengths, data point positions of *bb* (in grey), *bc* (in white) and other combinations (in black) are overlaid on the planes. (**A-B**) The upper row depicts the relative risk for psychiatric disease (here: MDD or ANX) in the younger subjects. Note a relatively low risk (<1, blue) for A1|A2 with similar lengths, including the *bb* and *bc* categories, a higher risk (>1, red) only for rare A1|A2 combinations, and a twisted shape of the central *bb*|*bc* area and the entire plane as indication of the A1-by-A2-interaction effect (**Supplementary Tab. 6**.**1**). (**C-D**) The lower row depicts the relative risk for psychiatric disease in the older subjects. Here, the A2 effect is not dependent on A1. Note the increase between *bb* and *bc*, as in the statistical analyses of these categories. **(A)** and **(C)** represent the planes without extrapolation of the model to non-represented values of A1 and A2, whereas **(B)** and **(D)** represent the estimated plane extrapolated to the entire data range. (**E**) represents the color scale denoting the relative risk, with yellow marking the turning point between reduced and increased disease risk.

**Figure 3.**
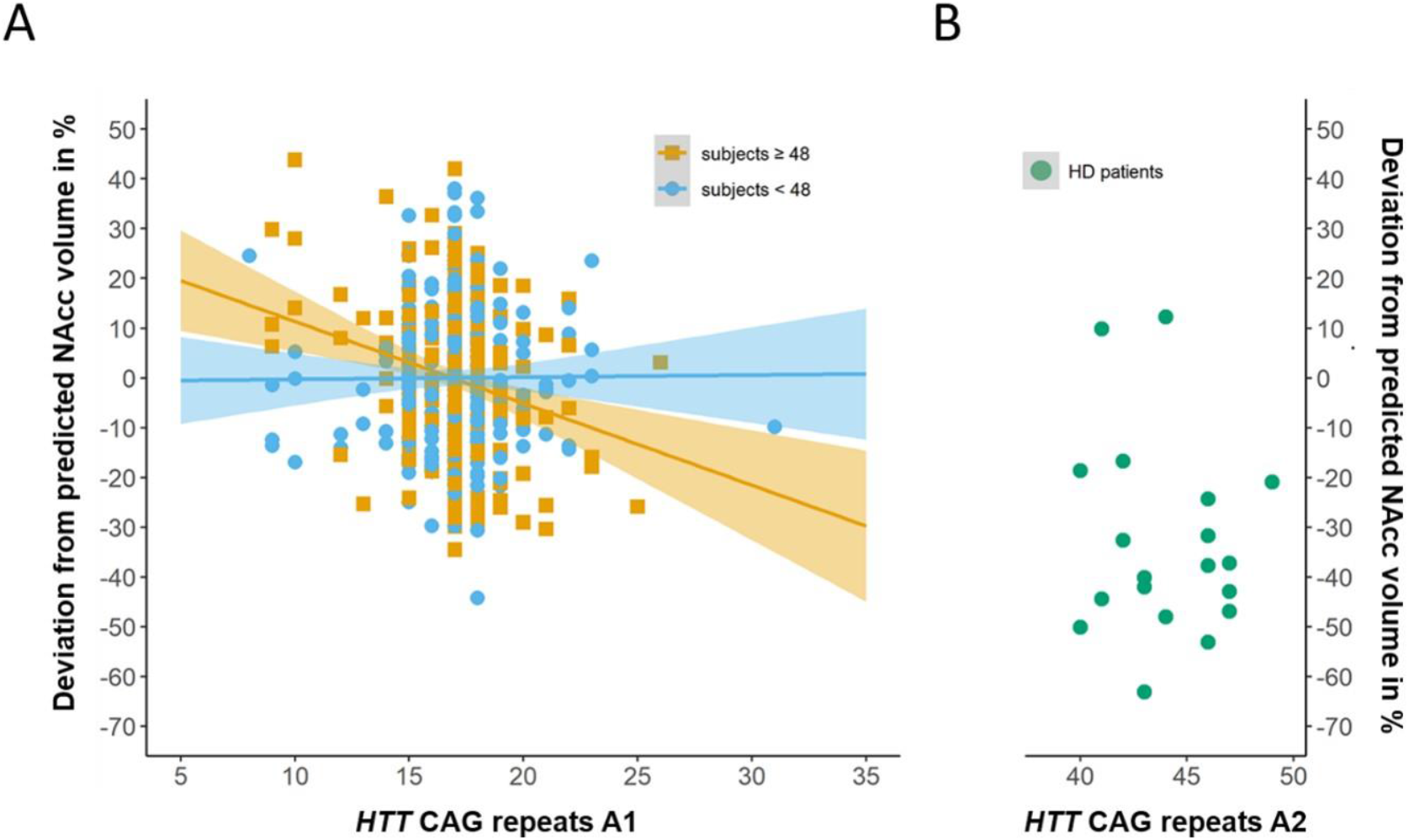
Interaction between age and *HTT* CAG repeat counts of A1 on accumbens volume. **(A)** Note negative slope between accumbens volume and *HTT* CAG repeat length on accumbens volume in subjects of 48 years and older (not separated by health and disease status). Units are percentage deviation between measured volume and the volume predicted from a model built on the MDD/CON sample using age, sex, estimated intracranial volume and coil type as predictors. **(B)** Green sample represents 19 patients with genetically proven HD for comparison. Their *HTT* CAG repeat counts on A1 were not considered in the analysis. Instead, their repeat counts on A2 are visualized on the x-axis.

#### HTT CAG repeats and MRI phenotypes

Effects of the *HTT CAG* repeat counts on the four subcortical volume measures, were investigated by multiple linear regression analyses using the predictors age, sex, case-control status, eICV, coil type, A2 and A1 (orthogonalized to A2 [A1_orth_] to avoid collinearity) and interaction terms A1_orth_-by-A2, age-by-A1_orth_ and age-by-A2. To not overlook nonlinear interaction effects, the effect of A1_orth_ was estimated for subsamples split at different A2 positions (**Supplementary Fig. 7**).

#### HTT CAG repeats and clinical variables

Clinical correlates of the *HTT* CAG repeat counts were explored by linear and logistic regression models for selected clinical variables (**Supplementary Table 10**) of the *MARS-Depressio*n Study using the predictors A1_orth_, A2, age (group; median split at 50 years), sex and interaction terms A1_orth_-by-A2, age-by-A1_orth_ and age-by-A2.

## RESULTS

### Distribution of *HTT* CAG repeat sizes

*HTT* CAG repeat sizes in 4195 subjects ranged from 7 to 39 repeats (**Fig. 1A**) (mean [SD] 18.4 [3.2]). Overall, five HD alleles in the reduced penetrance range (36-39 repeats)^13^ were identified, hereof three in CON (subject 1: female, 67 years, 38 *HTT* CAG repeats; subject 2: male, 61 years, 37 *HTT* CAG repeats; subject 3: male, 47 years, 39 *HTT* CAG repeats) and two in MDD patients (patient 1: female, 39 years, 38 *HTT* CAG repeats; patient 2: male, 24 years, 37 *HTT* CAG repeats). A total of 5.3% of all subjects (5.7% in the MDD sample, 5.3% in the ANX sample and 4.7% in the CON sample) had at least one allele in the range of 27-35. The most prevalent genotypes were *bb* and *bc* (63.8% and 24.3% respectively, **Fig. 1B** and **Supplementary Fig. 2**). The mean repeat length of A1 and A2 did not differ between patients and CON (**Supplementary Tab. 2**).

### Risk modulation of MDD and ANX by *HTT* CAG repeat sizes

#### Modelling CAG repeat sizes as categories

Patient status and allelic combinations of the predefined *HTT* CAG repeat ranges *a* (7-12), *b* (13-20), *c* (21-26), *d* (27-35) were only associated in subjects older than 48 years (Fisher’s exact test, p_FDR_=0.033; **Supplementary Tab. 5**). To better estimate effect sizes in these individuals, odds ratios (OR) were calculated for all allele combinations as long as a minimum of 5 subjects were available per *HTT* CAG repeat range and diagnosis: For *bb* (13-20|13-20) a lower risk for ANX was found (OR=0.463, CI=0.303-0.709, p_FDR_=0.002) whereas *bc* (13-20|21-26) was associated with an increased risk (OR=2.201, CI=1.408-3.440, p_FDR_=0.002) for ANX. Trends in the same direction were observed for MDD (*bb*: OR=0.807, CI=0.673-0.967, p_FDR_=0.051; *bc*: OR=1.282, CI=1.042-1.576, p_FDR_=0.051) (**Fig. 1C; Supplementary Tab. 6**). Robustness of these results was checked by varying the split point of repeat length between *b* and *c*, finding strongest ORs differences for MDD and ANX if the split point was set at 20, 21 or 22 (**Fig. 1D, Supplementary Tab. 7**). The effect direction, though, was the same for split points 19-23.

#### Modelling CAG repeat counts as continuous measures

In younger subjects, no significant effect for A1 or A2 or their quadratic extensions was detected, but a significant A1-by-A2 interaction (p=0.010) (**Supplementary Tab. 9**.**1**). Visually, this was reflected in three phenomena (**Fig. 2A–B**): First, a relatively low risk for the majority of allele combinations, second, a higher risk carried by rare A1|A2 combinations, and third, a ‘twisted’ shape of the plane indicating an influence of A2 mainly for low A1 values. In older subjects, a significant A2 effect (p=0.046) and a trend effect for its quadratic extension was detected, yet no A1-by-A2 interaction (**Supplementary Tab. 9**.**1**). Visually, this was reflected by a higher risk for larger A2 values across a broad A1 range [similar to the frequency difference between *bb*|*bc* in which one allele is kept stable in the *b* category (**Fig. 2C–D**)]. In the pooled sample with an age factor, A2 and age-by-A2 were significant, and trend effects were found for A2^2^-by-age and age-by-A1-by-A2 (**Supplementary Tab. 9**.**2**).

### Association between *HTT* CAG repeat counts and basal ganglia volumes

Effects of MDD status were detected for the pallidum and caudate, both showing volume increases in patients (**Tab. 1**). Exploratively, no interactions between *HTT* CAG repeat counts and MDD status were detected and thus the respective terms were removed from the final model. The lowest p-value was found for a negative age-by-A1_orth_ interaction effect on the nucleus accumbens (beta=-0.410, p=0.004, p_FDR_=0.017), followed by a trend positive correlation with A1_orth_ (beta=0.346, p=0.015, p_FDR_=0.058) and a nominally significant A1_orth_-by-A2-association (beta=-0.082, p=0.026, p_FDR_=0.104) (**Tab. 1**). **Fig. 3** depicts the age-by-A1_orth_ interaction effect: HD patients showed strong negative deviations from the expected volumes, yet the A1 correlation trajectory in the MCC|CON begins to overlap with the HD distribution.

**Table 1.** Association of basal ganglia volumes with *HTT* CAG repeat counts.

| Brain structure | N | Rs<sub>q</sub> | MDD status |  | A1<sub>orth</sub> |  | A2 |  | Age |  | A1<sub>orth</sub> × A2 |  | A1<sub>orth</sub> × Age |  | A2 × Age |  |
| --- | --- | --- | --- | --- | --- | --- | --- | --- | --- | --- | --- | --- | --- | --- | --- | --- |
|  |  |  | β | p<sub>FDR</sub> | β | p<sub>FDR</sub> | β | p<sub>FDR</sub> | β | p<sub>FDR</sub> | β | p<sub>FDR</sub> | β | p<sub>FDR</sub> | β | p<sub>FDR</sub> |
| Putamen | 544 | 0.475 | -0.020 | 0.651 | 0.226 | 0.178 | 0.114 | 0.554 | -0.458 | <0.001*** | -0.037 | 0.374 | -0.245 | 0.136 | -0.085 | 0.656 |
| Pallidum | 544 | 0.491 | -0.106 | 0.006* | -0.077 | 0.556 | -0.071 | 0.554 | -0.380 | <0.001*** | 0.009 | 0.792 | -0.006 | 0.962 | 0.082 | 0.656 |
| Caudate ncl. | 544 | 0.545 | -0.083 | 0.016* | 0.104 | 0.535 | -0.141 | 0.554 | -0.170 | <0.001*** | -0.057 | 0.154 | -0.090 | 0.625 | 0.115 | 0.656 |
| Ncl. accumbens | 544 | 0.404 | 0.016 | 0.651 | 0.346 | 0.058 | -0.091 | 0.554 | -0.490 | <0.001*** | -0.082 | 0.104 | -0.410 | 0.017* | 0.058 | 0.656 |

Note: N, number of available subjects; Rs<sub>q</sub>, adjusted R<sup>2</sup> of the full model; ncl., nucleus; β, standardized beta coefficient; p, p-value; \* and \*\*\* marking p-values < 0.05 and < 0.001, respectively; x, interaction; see methods section for full model description

### *HTT* CAG repeat sizes and clinical variables

Of all variables (**Supplementary Tab. 10**), we detected (nominally significant) positive correlations (p_FDR_>0.05) between A1_orth_ and apathetic syndrome at admission (p=0.044, p_FDR_=0.207) and at week 4 (p=0.009, p_FDR_=0.128), a negative correlation between A1_orth_ and remission (p=0.037, p_FDR_=0.207), and a positive correlation between A1_orth_-by-A2 and BDI-II at admission (p=0.009, p_FDR_=0.129) and HAMA at admission (p=0.047, p_FDR_=0.328)(**Tab. 2**). The effects of age on suicidal ideation and behavior and on a family history of MDD were not unexpected and in accordance with previous reports^40,41^. Following the *HTT* effects on the nucleus accumbens volume, we turned to specific BDI-II items reflecting anhedonia and variables reflecting addictive behaviour: Here, we found a correlation between anhedonia related BDI-II items at admission and A1_orth_-by-A2 (p=0.006) as opposed to a weaker effect for the complementary BDI-II items (p=0.019). No effects were found for cigarette smoking (**Supplementary Tab. 11**).

**Table 2.**
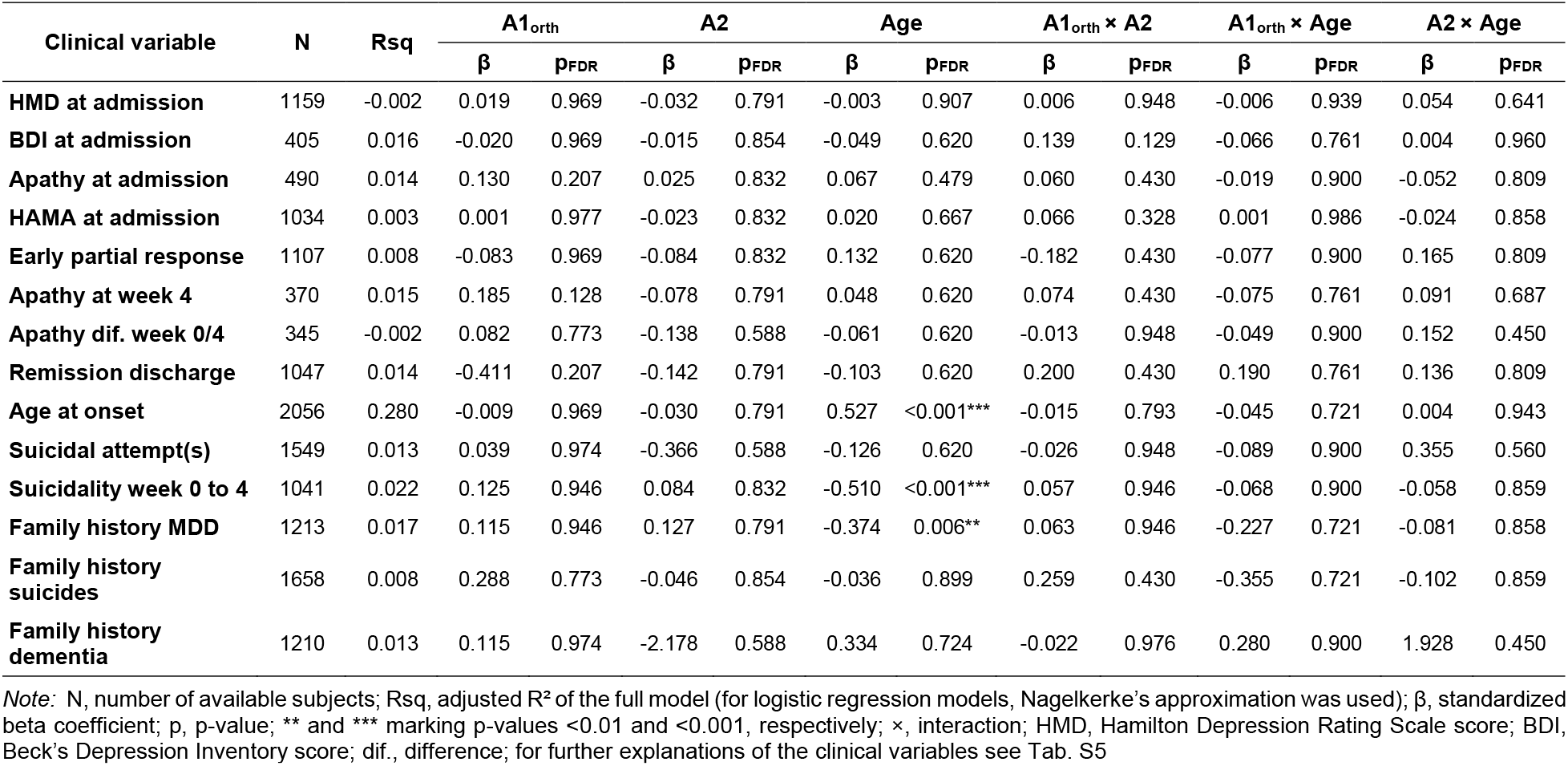
Association of clinical variables with *HTT* CAG repeat counts in MDD patients.

## DISCUSSION

We report on *HTT* CAG repeat variations in patients with MDD, ANX and psychiatrically healthy subjects of Caucasian origin. Beyond the diagnostic categories we also investigated dimensional psychiatric phenotypes and basal ganglia volumes, an established intermediate phenotype of premanifest HD.

### No overrepresentation of HD mutations in patients with MDD or ANX

*HTT* alleles carrying the HD mutation (>35 *HTT* CAG repeats) showed a frequency of ∼1/1315 in the patient group (MDD and ANX), and a frequency of ∼1/520 in CON. The latter was in line with a relatively high general frequency of HD mutations compared to registered HD cases in Western populations: Indeed, previous studies of HD mutations in general population samples in Canada, United States and Scotland, frequencies ∼1/400^12^, and ∼1/440 in Portugal^42^. The lack of over-representation of HD alleles in patients compared to CON was in contrast to a previous report^27^. For further clarification even larger sample sizes would need to be analyzed.

### HD alleles in the range of reduced penetrance or intergenerational instability

The three subjects of our psychiatrically healthy sample that were carriers of an HD allele in the range of reduced penetrance (36–39 CAG repeats) had already entered or passed mid-life (age range 47–67 years) and had no known diagnosis of HD, confirming the concept of reduced penetrance. Still, minor HD symptoms such as subtle motor, cognitive or behavioural abnormalities may have been missed as the original studies focused on psychiatric symptoms and complaints. While a positive family history of HD seems to be unfavourable^43^, there is no reliable empirical information about the penetrance risk of HD in *HTT* allele carriers in the reduced penetrance range. The range of 27-35 *HTT* CAG repeats was similarly represented in MDD, ANX and CON, matching reports in other western population samples of ∼3-6%^12,42,44^.

### Psychiatric disease risk modified by *HTT* CAG repeat ranges below 27

A further subdivision of *HTT* CAG repeats below 27 is not common practice in HD research as there is no known risk of genetic instability in this range and the length of the non-HD has not been shown to influence the motoric disease onset^45^. However, a variable lifetime risk modulation of depressive disorders by *HTT* CAG repeat counts below 27 has indeed been reported^11^. This encouraged us to introduce two further repeat cut-offs, resulting in three categories: *a* 7–12, *b* 13–20 and *c* 21–26 *HTT* CAG repeats.

As revealed by our frequency distribution analysis, ∼64% of the subjects had *HTT* CAG repeats in the range of *b* (13–20 repeats) on both alleles (i.e., *bb*), followed by ∼24% carrying the combination *bc* (13–20|21–26). We found that carrying the most frequently occurring *HTT* CAG repeat allele combination *bb* was associated with a lower risk of ANX in subjects ≥ 48 years whereas the allele combination *bc* was associated with an increased risk of ANX. Trends in the same directions were found for the risk of MDD. Two post-hoc analyses consolidated this result: First, we varied the definition of the boundaries between category *b* and category *c*, confirming that the risk change was indeed strongest around 21 repeats (**Fig. 1, Supplementary Fig. 4**). Second, given a slightly higher mean age of CON compared with ANX, we excluded an influence of age stratification on our risk results by respective age comparisons between *bb* and *bc* in CON, finding no such bias. In summary of these frequency analyses we extend the current literature on the role of huntingtin for psychiatric disease risk^11,27,28,46^ that a critical threshold of ∼21 CAG repeat counts of the longer allele might play a role for psychiatric risk, yet depending on age.

### Dimensional analysis of A1 and A2 under consideration of their intrinsic correlation

The mean *HTT* CAG repeat count of all quantified alleles in our study lay within the expected range between 18.4 and 18.7 as reported for subjects of European descent^1^. So far, the correlation of the ‘longer’ and ‘shorter’ *HTT* allele has been interpreted as evidence for assortative mating - a mating pattern based on similarities^47,48^. Yet, as the ‘shorter’ allele, by definition, represents the lower boundary for the ‘longer’, and as soon as this definition is lifted, e.g., by permutation analysis with random assignment to two groups, this correlation disappears (**Supplementary Fig. 3B**), likely not justifying to conclude evidence for assortative mating.

We then studied risk effects independently of pre-defined CAG repeat boundaries, confirming that key role of age: While in older subjects the risk was positively correlated with A2 (with *no* A1-by-A2 interaction), it was modulated by an A1-by-A2 interaction in younger subjects, driven by strongly discrepant values of A1 and A2. The relatively lowest psychiatric risk was indeed found for allele combinations with similar *HTT* CAG repeat counts that included the *bb* and *bc* categories. Post-hoc, the categorial allele combinations were also analyzed in the younger subjects, revealing a trend risk effect of *bb* and trend protective effect of *bc* for ANX (**Supplementary Fig. 5**), inversely to the older subjects. From the synopsis of the categorical and dimensional analyses we thus conclude that ageing itself – by a still unknown (patho-)physiological mechanisms– seems to lead to a reversal of the risk effect of genotype combinations *bb* and *bc*. Our data also convey that the shorter allele should not be neglected in the exploration of the function of *HTT* for brain circuits.

### MRI basal ganglia correlates of *HTT* CAG repeat variation

Focusing on four basal ganglia markers predominantly affected in HD, we detected a negative age-by-A1 effect on the nucleus accumbens volume. This effect was independent of the MDD status, and MDD status itself was not associated with nucleus accumbens volume, in line with a large meta-analysis^49^. There is a plenitude of structural MRI studies deciphering the volumetric effects of *HTT* CAG repeats on the brain in prodromal and manifest HD^16,18,50–53^ as well as on atrophy progression^22,54^. We found no direct correlations between A2 and basal ganglia markes, yet, for the nucleus accumbens, a longer A2 intensified the effect of A1, with a critical threshold of about 20-21 CAG repeat counts of A2 (**Supplementary Fig. 7**). Whereas for children and adolescents, a positive correlation between A2 and total grey matter has been reported specifically for females^55^, for adults, our report is the first on associations between the *lower* count *HTT* CAG repeat allele and basal ganglia markers in the absence of an HD allele. Involvement of the nucleus accumbens in cognitive decline and dementia has been highlighted in a population-based study^56^, and neuropathological involvement of the nucleus accumbens in HD was found associated with psychiatric rather than motor symptoms at disease onset^23^, which conforms to the weak but still detectable correlations of the repeat counts with depression and anxiety levels. We cannot pin down the mechanisms underlying the vulnerability of the nucleus accumbens towards huntingtin dependent aging processes. Several cellular processes that are modulated also by wildtype *HTT*^*57,58*^ (e.g., transcription of BDNF in cortical neurons, axonal transport of neurotrophic substances to the striatum, regulation of tissue maintenance in a wider sense) are plausible.

### *HTT* CAG repeat sizes and depression phenotypes

In order to analyze the influence of *HTT* CAG repeat sizes on the depression phenotype at a fine-grain level, we analyzed more fine-granular MDD variables^23^ covering acute symptoms, treatment response, individual depression history and family history. In brief, no robust associations with either allele or their interaction emerged. The relatively strongest effect was a negative correlation with age at MDD onset which replicates the observation that HD patients with a psychiatric onset tend to be younger than those with a motor onset^23^. Our results regarding the nucleus accumbens led us to explore anhedonia-related BDI items and substance abuse, both potentially associated disturbed dopaminergic signaling: We found that indeed the anhedonia items correlated with an A1-by-A2-interaction term slightly stronger than the remaining items. The direction of the interaction effect was positive, aligning with the inverse direction of the same term regarding the nucleus accumbens volume. No effect on cigarette consumption as a more defined addiction behaviour was detected. Finally, as we observed a more pronounced risk modulation by *HTT* CAG repeats for ANX than MDD, we explored the subtype of anxious depression, but detected no change in MDD risk (**Fig. 1, Supplementary Fig. 8**). Overall, we found no strong associations of *HTT* CAT repeat variations with specific clinical MDD or ANX profiles.

## Conclusion

We conclude that *HTT* CAG repeats in the most frequently occurring ranges modulate the risk of depressive and anxiety disorders, yet only in subjects older than 48 years. This risk was unfavourably modulated by higher CAG repeat counts below the HD cut-off with a non-linear risk increase around 21 repeats in at least one allele. In younger subjects, this effect was absent or even trending towards a protective effect. We also observed that with higher age the nucleus accumbens volume was negatively correlated with *HTT* CAG repeat sizes of the shorter allele. Our findings corroborate the potential of *HTT* CAG repeats in the non-HD range to exert age-dependent modulating effects on the susceptibility towards common psychiatric diseases and basal ganglia structure.

## Acknowledgements

We thank Monika Rex-Haffner and Laura Diener for excellent technical assistance and Marcus Ising for valuable information and discussions. We thank Andreas Bender and Dorothee P. Auer for patient recruitment and acquisition of the Huntington’s Disease MRI study. We are further grateful to Rosa Schirmer for great support of data management, and we thank all study participants for their support of clinical studies at the MPIP.

