## Supplementary Information for "Huntingtin CAG repeat size variations outside the Huntington’s disease complex: associations with depression and anxiety phenotypes and basal ganglia structure"

### SUPPLEMENTARY DATA

### SUPPLEMENTARY FIGURES

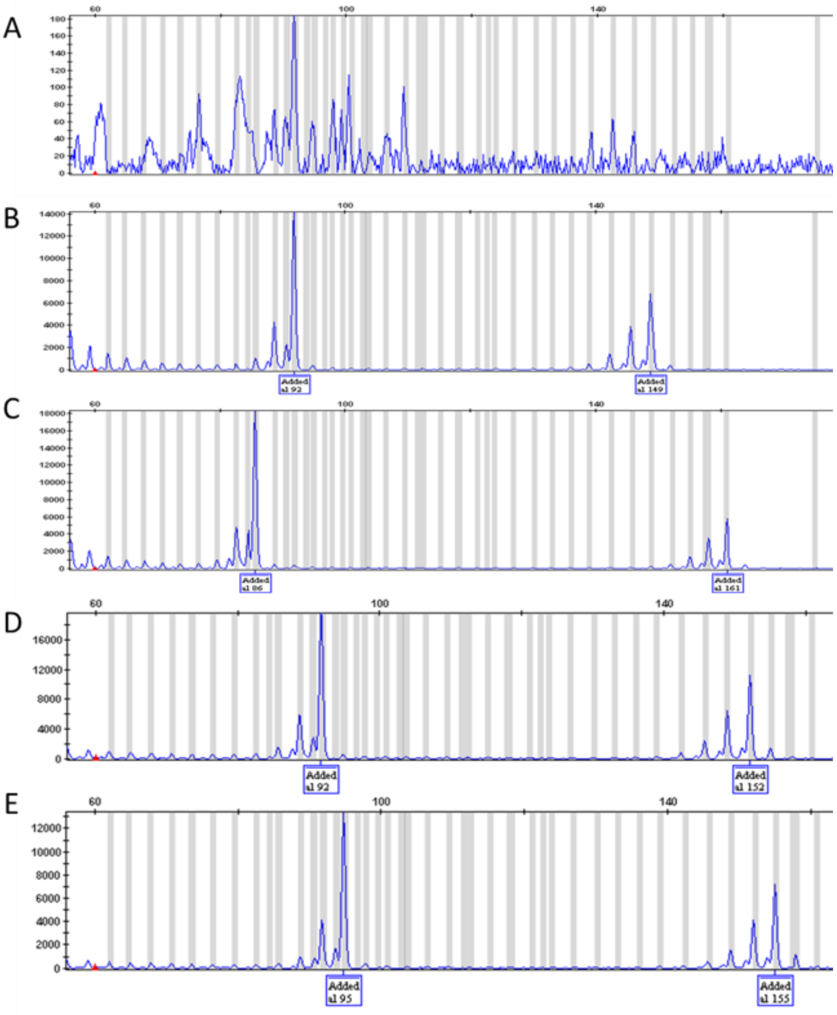


**Supplementary Figure 1. Selected GeneMapper screen shots of the fragment analysis.** The y-axis represents relative fluorescent units, the x-axis represents base pair sizes as estimated from an internal size standard. **(A)** Negative control (water). **(B)** Standard B from the National Institute of Standards & Technology, USA, *HTT*-CAG-repeat triplets 17/36. **(C)** Standard C from the National Institute of Standards & Technology, USA, *HTT* CAG repeat triplets 15/40. **(D)** Study subject with converted *HTT* CAG repeat triplets 17/37. **(E)** Study subject with converted *HTT* CAG repeat triplets 18/38.

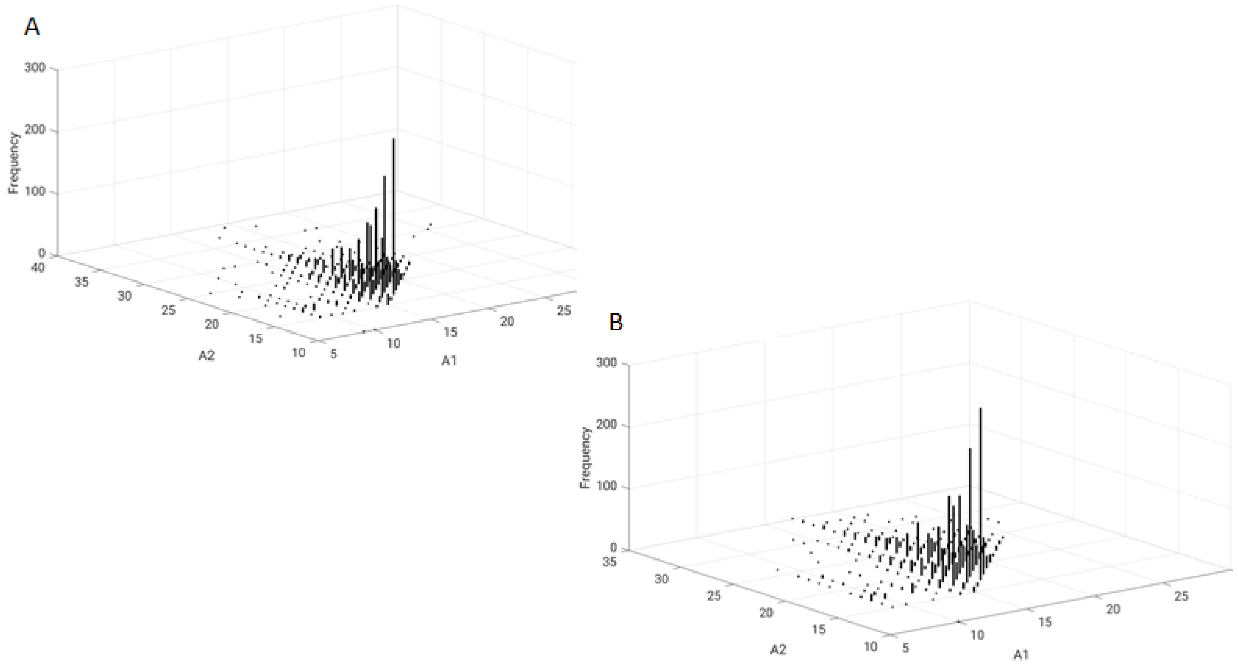


**Supplementary Figure 2. Illustration of the frequency of all A1|A2 pairs.** 3D-histograms were plotted to illustrate the density of the data source for all A1|A2 pairs for subjects < 48 years **(A)** and subjects of ≥ 48 years **(B)**. The x and y-axis represent *HTT* CAG repeat counts of allele A1 or A1 as indicated and the z-axis the frequency.

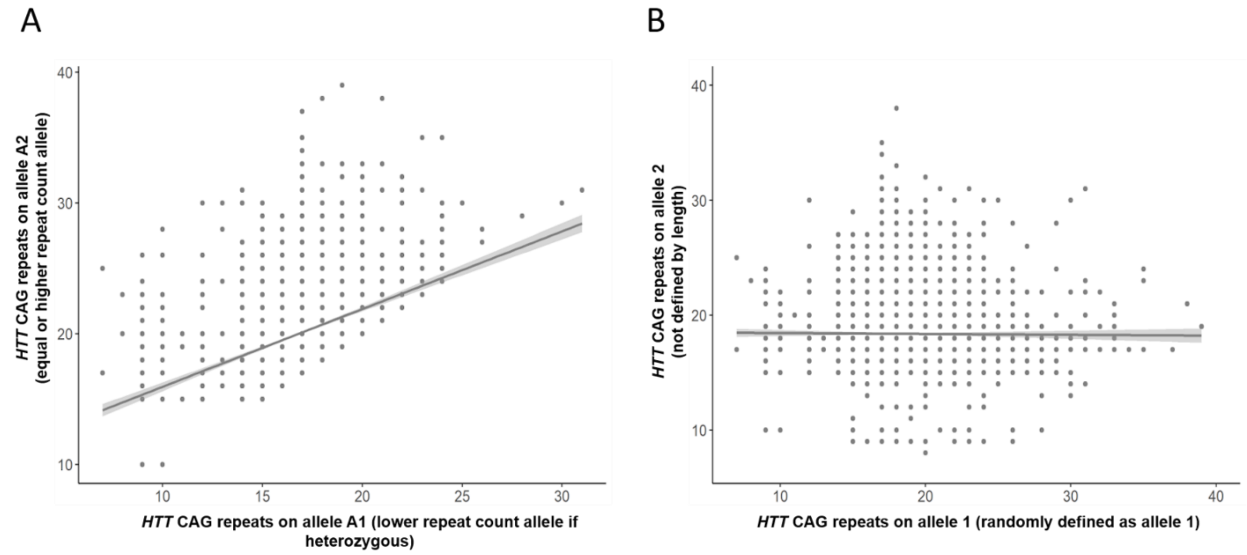


**Supplementary Figure 3. Testing for correlation between *HTT* CAG repeat counts of both alleles.** *HTT* CAG repeats of both alleles were tested for correlations. We previously defined two *HTT* allele types: A1 defined as lower *HTT* CAG repeat count allele and A2 as homozygous or higher *HTT* CAG repeat count allele. **(A)** shows the pseudocorrelation resulting from the presorting of *HTT* alleles by repeat count, x-axis A1 and y-axis A2. **(B)** Shows no correlation after subject-wise attributing repeat counts randomly to two categories not defined by length (allele1 x-axis and allele 2 y-axis), (r=-0.01, p=0.62).

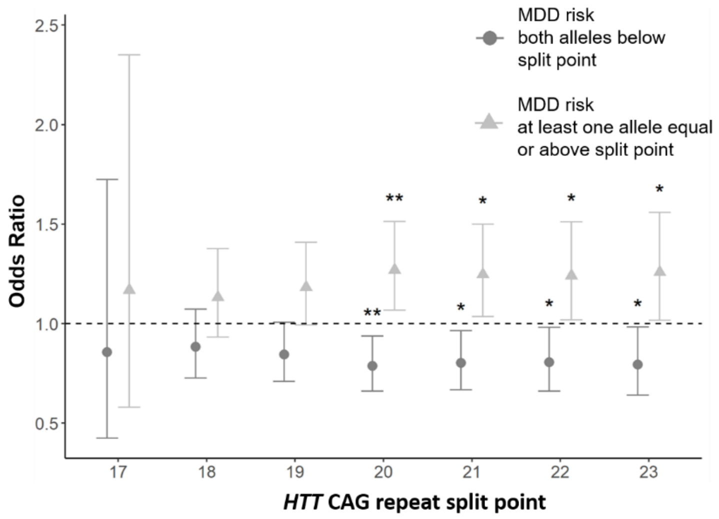


**Supplementary Figure 4. Association between *HTT* CAG repeat sizes and risk of MDD in subjects ≥ 48 years.** ORs were calculated to evaluate MDD risk for different allele combinations in subjects ≥ 48 years. Error bars indicate confidence intervals. ORs are shown at different split points comparing MDD risk of the group having both alleles below the split point to the group having at least one allele of the length of the split point or larger. * and ** marking nominal p-values < 0.05 and 0.01 respectively. Effects were not statistically significant after correction for multiple comparisons.

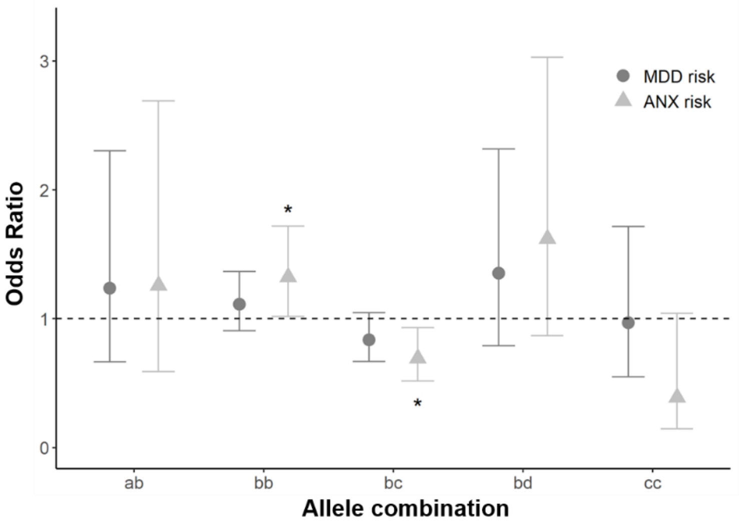


**Supplementary Figure 5. Association between *HTT* CAG repeat sizes and risk of MDD or ANX in subjects < 48 years.** To evaluate the risk of MDD or ANX for allele combinations *ab, bb, bc, bd and cc* in subjects younger than 48 years ORs were calculated. Error bars indicate confidence intervals. Nominal significant p-values are indicated, * marking nominal p-values < 0.05. Effects were not statistically significant after correction for multiple comparisons.

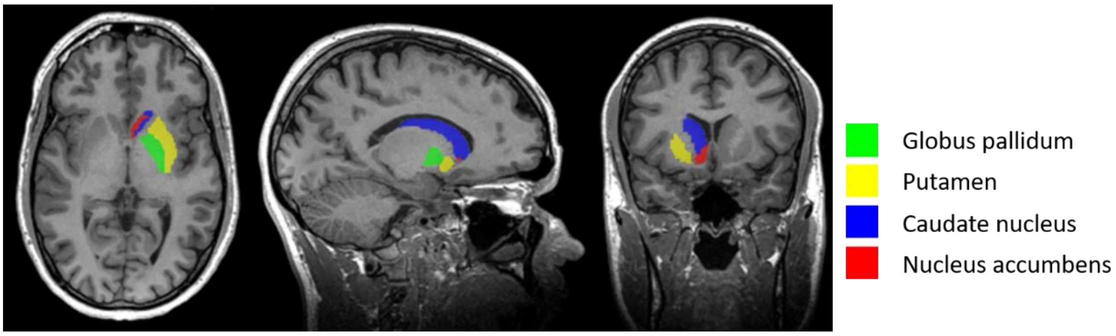


**Supplementary Figure 6. Segmentation of selected subcortical volumes.** Segmented unilateral pallidum, putamen, caudate nucleus and nucleus accumbens (FreeSurfer version 5.3) of an exemplary case are shown.

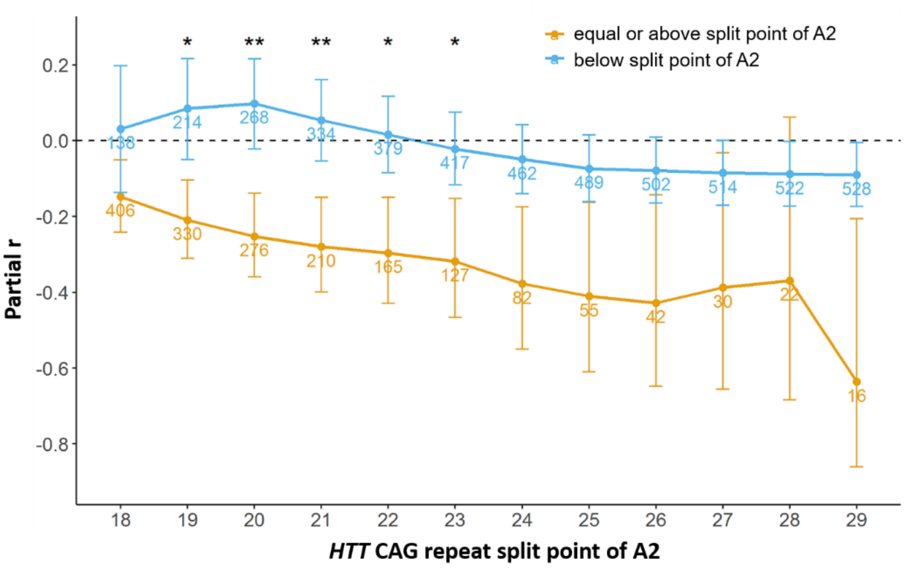


**Supplementary Figure 7. Association between *HTT* CAG repeat counts of A1 and nucleus accumbens volume depending on A2 split point.** Subjects independent of disease status were analyzed. Note negative partial correlation values for all associations built for subjects equal or above the A2 split point, and no correlation or slightly positive correlation for subjects below the A2 split point. Asterisks indicate nominally significant interaction effects (A1 length by A2 split): * $p < 0.05$, ** $p < 0.001$.

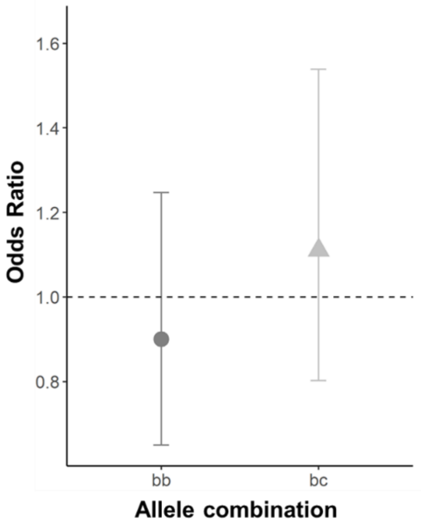


**Supplementary Figure 8. ORs calculated for a subsample of MDD with HAMA at admission scores of at least 23.** To evaluate the MDD risk of subjects of 48 years or older and HAMA at admission scores equal or above 23 OR were calculated using the age matched CON group. The threshold of 23 was chosen as it represents the median HAMA score in the MDD group. Error bars indicate confidence intervals.

### SUPPLEMENTARY METHODS

#### Subjects

The Recurrent Unipolar Depression (RUD) study and the Munich Antidepressant Response Signature (MARS) project and their corresponding control groups were the sources for subjects included in the present study. The MARS project comprised two substudies: MARS Depression and MARS Anxiety. For subsamples of the RUD study and the MARS project, structural MRI data were available as detailed below.

*RUD Study*

The RUD study was a cross sectional case-control study comprising patients with recurrent unipolar depression and control subjects. Patients were recruited at the clinic of the Max Planck Institute of Psychiatry (MPIP), Munich, Germany, and two further psychiatric hospitals located in southern Bavaria. Information about demographics, ethnicity, family, individual and medical history were acquired using a questionnaire completed by the participants. All patients were assessed by the semi-structured interview Schedule for Clinical Assessment in Neuropsychiatry (SCAN)[1]. Recurrent unipolar depression was diagnosed according to the International Statistical Classification of Diseases and Related Health Problems (ICD-10)[2] and/or the Diagnostic and Statistical Manual of Mental Disorders IV (DSM-IV)[3]. Caucasians with at least two moderate-to-severe depressive episodes were included. Exclusion criteria comprised (i) the presence of manic or hypomanic episodes, (ii) mood incongruent psychotic symptoms, (iii) a lifetime diagnosis of intravenous drug abuse, (iv) depressive symptoms only secondary to alcohol/substance abuse/dependence or (v) depressive symptoms only secondary to a medical illness or medication and (vi) age younger than 18 years.

Control subjects matched for ethnicity, sex and age were recruited at the MPIP. They were selected from a Munich-based community sample. Only individuals negative for depression and anxiety disorders as assessed by a SCAN interview were included. The study was approved by the responsible Ethics Committee (Bayerische Landesärztekammer) and written informed consent was obtained from all subjects. The RUD study sample has been described and analyzed before in several genetic association studies[4-6]. In the present study 904 cases and 1028 controls from the RUD study were included.

*MARS-Depression*

This study was a prospective multi-center naturalistic observational study of treatment outcomes in acutely depressed in-patients with different types of depressive disorders, mainly MDD, but also bipolar disorder (see overview[7]). The study aimed at generating a large databank of sociodemographic, psychopathological, and biological data along with longitudinal psychometry with weekly ratings (e. g. Hamilton Depression Rating Scale 21 items version [HMD][8], Beck Depression Inventory-II [BDI][9] or ratings based on the Association for Methodology and Documentation in Psychiatry [AMDP] system). For our clinical correlation analyses in the present study, we focused on a defined subset of clinical items appropriate in the given context. This selection was based on (i) its reported relatedness to the HD neuropsychiatric symptom spectrum and (ii) to its broad availability across the sample. Hence, we selected variables in the domain of acute symptoms of depression and anxiety (HMD and Hamilton Anxiety Rating Scale [HAMA] total scores, apathetic syndrome as derived from the AMDP rating system), depression treatment response (HMD at week 2, remission at discharge), depression history (suicide attempts, age at onset), family history of major depressive disorder, suicides and dementia) and addictive behavior (cigarettes per day). Regarding alcohol use disorders, the respective phenotypes were too scarce due to the studies' inclusion criteria or heterogeneously measured in the different source studies to perform a reliable analysis.

Study subjects were recruited at the MPIP in Munich and collaborating hospitals in southern Bavaria or Switzerland. Only Caucasians were included. Study inclusion was performed within the first days after admission. ICD-10 diagnoses were obtained by trained psychiatrists. General exclusion criteria were depressive syndromes secondary to any medical or neurological condition, the presence of manic, hypomanic or mixed affective symptoms, lifetime diagnosis of alcohol dependence, illicit drug abuse or the presence of severe medical conditions. In the present study we included only patients from the MARS Depression sample with MDD as main diagnosis (n=1235) selected by the ICD-10 codes F32 (Major depressive disorder, single episode) or F33 (Major depressive disorder, recurrent episodes), or patients with an anxiety disorder (n=5) selected by the ICD-10 codes F40 (Phobic anxiety disorders) or F41 (Other anxiety disorders) as main diagnosis. Patients with bipolar disorder were excluded.

*MARS-Anxiety*

Patients consecutively admitted to the Anxiety Disorders Outpatient Clinic at the MPIP for diagnosis and treatment of an anxiety disorder were recruited. Patients were diagnosed according to ICD-10 and DSM-IV by trained psychiatrists and examined as described before[10,11]. Patients with anxiety disorders due to a medical or neurological condition or a comorbid axis II disorder were excluded. All patients were Caucasians. We included 490 patients in the present study that met the diagnostic criteria for an anxiety diagnosis according to ICD-10 code F40 (Phobic anxiety disorders) or F41 (Other anxiety disorders). From this sample we also included 9 patients that had MDD as the main diagnosis to be analyzed within the combined MDD sample.

*MARS-Controls*

MARS control subjects were randomly selected from the Munich resident's registry. The subjects were screened for the presence of anxiety and affective disorders and were only included if their screening result was negative. Control subjects were matched with the MARS Depression patients for age, sex and ethnicity. 541 controls were eligible for our present study.

The MARS project comprising the sub-studies MARS-Depression, MARS-Anxiety and MARS-Controls, was approved by the local Ethics Committee of the Ludwig Maximilian University, Munich, Germany. Written informed consent was obtained from all participants after the study protocol and all potential risks have been explained in detail.

*Molecular methods*

The following PCR conditions were applied: 80 ng of genomic DNA, 1.25 mmol $MgCl_2$, 1 x Buffer AmpliTaq Gold 360 (Applied Biosystems), 0.05 U/µl AmpliTaq Gold 360 DNA Polymerase (Applied Biosystems), 0.125 µmol each primer, 0.25 mmol deoxyribonucleotide triphosphate and 12% dimethyl sulfoxide in a 10 µl reaction. The following cycling conditions were used: initial denaturation at 94°C for 4 minutes, 35 cycles of denaturation at 94°C for 30 seconds, annealing at 65°C for 30 seconds, extension at 72°C for 45 seconds, and one final extension of 15 minutes at 72°C. Every PCR included a negative control without genomic DNA and DNA with predetermined *HTT* CAG repeat allele sizes. All assessments were done with cases and controls randomized on 96- or 384-well plates.

*Magnetic resonance imaging (MRI) samples and processing*

MRI data were subsamples of the *MARS-Depression/-Controls*[13], and the RUD Study[14] as reported before in imaging genetics and morphometry studies[12,15]. Overall, 332 MDD patients (224 *MARS-Depression*, 108 RUD Study), and 212 CON (21 *MARS-Controls*, 191 RUD Study) were included. To validate the segmentation procedure and as reference for effect sizes, a sample of 20 HD patients was also analyzed[16]. Volumetric measures were generated by the

FreeSurfer algorithm[17,18] as applied to high resolution T1-weighted images[12,16]. Segmentation quality control followed standardized protocols (http://enigma.ini.usc.edu/protocols/imaging-protocols). Analysis of covariance comparing HD with healthy controls (correcting for age, sex, estimated intracranial volume [eICV] and coil type) confirmed volumetric deficits in the putamen and pallidum (~43%), nucleus accumbens (~36%) and caudate nucleus (~23%), and no hippocampal or amygdala deficits, in line with reports on (pre-symptomatic) HD using the same technique[19]. As intermediate phenotypes for the *HTT* allele status we focused on these four bilateral subcortical volumes (**Supplementary Fig. 6)**.

### SUPPLEMENTARY RESULTS

*Modelling CAG repeat sizes as categories*

When comparing ORs for MDD between subjects with both alleles below the split point against ones having at least one allele equal or above the split point (**Supplementary Fig. 4**, **Supplementary Tab. 8**) results similar to those seen in **Figure 1C** and **D** were found. Last, given a 2-4 years higher average age of CON compared with ANX for the allele combinations *bb* and *bc* (**Supplementary Tab. 4.1**), we probed if demographic differences between these combinations are present in CON: No such differences were detected in CON or the patient groups (**Supplementary Tab 4.2**), and no correlations between the repeat counts and age or sex in CON (**Supplementary Tab. 1**) were detected.

*Association between HTT CAG repeat counts and basal ganglia volumes*

To not overlook nonlinear interaction effects on the nucleus accumbens, the effect of $A1_{orth}$ was estimated for different A2 split points (**Supplementary Fig. 7**), showing negative correlations with $A1_{orth}$ in the subgroup with A2 *above* the split point, and no or positive correlations in the subgroup with A2 *below* the split point. This pattern was stable over a large range of split points and most pronounced split points 20 and 21. In summary, no direct A2 effect on the nucleus accumbens was found, yet, A2 *modulated* the association with A1.

*HTT CAG repeat sizes and clinical variables*

In consideration of the detected association of *HTT* CAG repeats with ANX in older patients, we recalculated ORs for the MDD subgroup of *anxious depression* (MDD patients with a HAMA score equal or above 23 (median) at admission, finding no effect of higher anxiety levels in MDD (*bb:* OR=0.90; CI=0.65-1.25; *bc:* OR=1.11; CI=0.80-1.54; p=0.53; see **Supplementary Fig. 8**).

### SUPPLEMENTARY TABLES

#### Supplementary Table 1. Association of age and sex with *HTT* CAG repeat counts within CON, MDD and ANX

| | CON | | | | MDD | | | | ANX | | | |
|---|---|---|---|---|---|---|---|---|---|---|---|---|
| | age | | sex | | age | | sex | | age | | sex | |
| **Alleles** | **r** | **p** | **t** | **p** | **r** | **p** | **t** | **p** | **r** | **p** | **t** | **p** |
| **A1** | 0.034 | 0.179 | 1.818 | 0.069 | -0.002 | 0.917 | 0.818 | 0.414 | 0.015 | 0.740 | -1.468 | 0.143 |
| **A2** | 0.001 | 0.967 | 0.878 | 0.380 | 0.011 | 0.597 | -0.667 | 0.505 | 0.089 | 0.049* | 0.067 | 0.947 |

*Note:* A1, lower *HTT* CAG repeat count on one allele; A2 higher or equal repeat count on other allele; r, Pearson correlation coefficient; t, t-statistic; p, p-value; * marking p-values < 0.05

**Supplementary Table 2. Comparison of mean *HTT* CAG repeat counts between groups MDD, ANX and CON**

| Alleles | MDD (N=2136) | ANX (N=493) | CON (N=1566) | df | F | p |
|---|---|---|---|---|---|---|
| **A1** | 16.9 ± 2.0 | 16.6 ± 2.1 | 16.8 ± 2.1 | 2 | 2.2 | 0.11 |
| **A2** | 20.1 ± 3.4 | 19.8 ± 3.2 | 20.0 ± 3.4 | 2 | 1.2 | 0.31 |

*Note:* A1, lower *HTT* CAG repeat count on one allele; A2 higher or equal repeat count on other allele; mean values ± standard deviation; df, degrees of freedom; F, F-statistics of main effect (ANOVA) of diagnosis (3-level factor); p, p-value

**Supplementary Table 3. Comparison of mean *HTT* CAG repeat counts between groups MDD, ANX and CON corrected for age and sex**

| | A1 | | A2 | |
|---|---|---|---|---|
| **Variable** | **F** | **p** | **F** | **p** |
| **Diagnosis** | 1.719 | 0.179 | 0.870 | 0.419 |
| **Age** | 0.626 | 0.429 | 0.958 | 0.328 |
| **Sex** | 1.309 | 0.253 | 0 | 0.994 |

*Note:* A1, lower *HTT* CAG repeat count on one allele; A2 higher or equal repeat count on other allele; F, F-statistics of main effect; p, p-value

**Supplementary Table 4.1. Mean ages and prevalence of sexes compared between diagnostic groups for allele combinations *bb* and *bc***

| | | < 48 years | | | | | | ≥ 48 years | | | | | |
|---|---|---|---|---|---|---|---|---|---|---|---|---|---|
| | | ***bb*** | | | ***bc*** | | | ***bb*** | | | ***bc*** | | |
| | | **CON** | **MDD** | **ANX** | **CON** | **MDD** | **ANX** | **CON** | **MDD** | **ANX** | **CON** | **MDD** | **ANX** |
| **age in years (mean±sd)** | | 36.5 ±7.75 | 36.6 ±7.95 | 32.8 ±7.91 | 36.9 ±7.14 | 36.1 ±8.32 | 34.0 ±8.32 | 59.2 ±7.93 | 59.5 ±8.40 | 56.7 ±7.79 | 58.8 ±8.02 | 59.5 ±8.54 | 55.4 ±6.27 |
| **n female (%)** | | 249 (60.7) | 347 (57.2) | 144 (53.1) | 119 (65.0) | 116 (50.9) | 49 (59.0) | 393 (65.1) | 445 (60.2) | 35 (76.1) | 116 (61.4) | 187 (61.9) | 21 (60.0) |
| **n male (%)** | | 161 (39.3) | 260 (42.8) | 127 (46.9) | 64 (35.0) | 112 (49.1) | 34 (41.0) | 211 (34.9) | 294 (39.8) | 11 (23.9) | 73 (38.6) | 115 (38.1) | 14 (40.0) |
| | | | **MDD vs CON** | **ANX vs CON** | | **MDD vs CON** | **ANX vs CON** | | **MDD vs CON** | **ANX vs CON** | | **MDD vs CON** | **ANX vs CON** |
| **age** | **t** | | 0.209 | -5.979 | | -1.123 | -2.752 | | 0.534 | -2.085 | | 0.873 | -2.860 |
| | **p** | | 0.834 | < 0.001*** | | 0.262 | 0.007** | | 0.593 | 0.042* | | 0.383 | 0.006* |
| **sex** | **$Chi^2$** | | 1.140 | 3.552 | | 7.734 | 0.642 | | 3.128 | 1.845 | | 0.001 | 0.000 |
| | **p** | | 0.286 | 0.059 | | 0.005** | 0.423 | | 0.077 | 0.174 | | 0.980 | 1.000 |

*Note*: *b*, *HTT* CAG repeat range 13-20; *c*, *HTT* CAG repeat range 21-26; *bb* and *bc*, allelic combinations of ranges *a* and *b*; sd, standard deviation; n, number; t, t-statistic; p, p-value; *, ** and *** marking nominal p-values < 0.05; < 0.01; < 0.001, respectively

**Supplementary Table 4.2. Mean ages and prevalence of sexes compared between allele combinations *bb* and *bc* for diagnostic subgroups**

| | CON age (mean±sd) | | | | MDD age (mean±sd) | | | | ANX age (mean±sd) | | | |
|---|---|---|---|---|---|---|---|---|---|---|---|---|
| | ***bb*** | ***bc*** | **t** | **p** | ***bb*** | ***bc*** | **t** | **p** | ***bb*** | ***bc*** | **t** | p |
| **< 48 years** | 36.5 ±7.75 | 36.9 ±7.14 | -0.746 | 0.456 | 36.6 ±7.95 | 36.1 ±8.32 | 0.744 | 0.457 | 32.8 ±7.91 | 34.0 ±8.32 | -1.221 | 0.227 |
| **≥ 48 years** | 59.2 ±7.93 | 58.8 ±8.02 | 0.594 | 0.553 | 59.5 ±8.40 | 59.5 ±8.54 | -0.054 | 0.957 | 56.7 ±7.79 | 55.4 ±6.27 | 0.875 | 0.384 |

| | CON n female (%) | | | | MDD n female (%) | | | | ANX n female (%) | | | |
|---|---|---|---|---|---|---|---|---|---|---|---|---|
| | ***bb*** | ***bc*** | **Chi²** | **p** | ***bb*** | ***bc*** | **Chi²** | **p** | ***bb*** | ***bc*** | **Chi²** | p |
| **< 48 years** | 249 (60.7) | 119 (65.0) | 0.818 | 0.366 | 347 (57.2) | 116 (50.9) | 2.405 | 0.121 | 144 (53.1) | 49 (59.0) | 0.670 | 0.413 |
| **≥ 48 years** | 393 (65.1) | 116 (61.4) | 0.700 | 0.403 | 445 (60.2) | 187 (61.9) | 0.194 | 0.659 | 35 (76.1) | 21 (60.0) | 1.716 | 0.190 |

*Note*: *b*, *HTT* CAG repeat range 13-20; *c*, *HTT* CAG repeat range 21-26; *bb* and *bc*, allelic combinations of ranges *a* and *b*; sd, standard deviation; n, number; t, t-statistic; p, p-value

**Supplementary Table 5. Comparison of prevalence of *HTT* allelic combinations between MDD patients, ANX patients and CON for entire sample and split by median age (48 years)**

| Allele combination | All ages | | | | < 48 years | | | | ≥ 48 years | | | |
|---|---|---|---|---|---|---|---|---|---|---|---|---|
| | MDD | ANX | CON | $p_{FDR}$ | MDD | ANX | CON | $p_{FDR}$ | MDD | ANX | CON | $p_{FDR}$ |
| | N | N | N | | N | N | N | | N | N | N | |
| **aa** | 2 | 0 | 1 | 0.314 | 1 | 0 | 1 | 0.314 | 1 | 0 | 0 | 0.033* |
| **ab** | 58 | 17 | 39 | | 28 | 12 | 16 | | 30 | 5 | 23 | |
| **ac** | 13 | 7 | 12 | | 5 | 5 | 4 | | 8 | 2 | 8 | |
| **ad** | 0 | 0 | 2 | | 0 | 0 | 2 | | 0 | 0 | 0 | |
| **bb** | 1346 | 317 | 1014 | | 607 | 271 | 410 | | 739 | 46 | 604 | |
| **bc** | 530 | 118 | 372 | | 228 | 83 | 183 | | 302 | 35 | 189 | |
| **bd** | 108 | 24 | 58 | | 40 | 20 | 21 | | 68 | 4 | 37 | |
| **be** | 1 | 0 | 3 | | 1 | 0 | 1 | | 0 | 0 | 2 | |
| **cc** | 63 | 8 | 52 | | 29 | 5 | 21 | | 34 | 3 | 31 | |
| **cd** | 13 | 1 | 12 | | 4 | 1 | 4 | | 9 | 0 | 8 | |
| **ce** | 1 | 0 | 0 | | 1 | 0 | 0 | | 0 | 0 | 0 | |
| **dd** | 1 | 1 | 1 | | 1 | 1 | 1 | | 0 | 0 | 0 | |

*Note:* a, *HTT* CAG repeat range 7-12; b, *HTT* CAG repeat range 13-20; c, *HTT* CAG repeat range 21-26; d, *HTT* CAG repeat range 27–35; e, > 35 *HTT* CAG repeats; N, number of available subjects; p-values as calculated by Fisher's exact test, * marking p-values < 0.05

**Supplementary Table 6. Association of disease risk with allelic combinations of predefined *HTT* CAG repeat ranges (≥ 48 years)**

| Genotype | MDD | | | ANX | | |
|---|---|---|---|---|---|---|
| | OR | CI | $p_{FDR}$ | OR | CI | $p_{FDR}$ |
| **ab** | 0.988 | 0.570–1.712 | 0.964 | 2.123 | 0.788–5.721 | 0.213 |
| **ac** | 0.756 | 0.283–2.021 | 0.720 | - | - | - |
| **bb** | 0.807 | 0.673–0.967 | 0.051 | 0.463 | 0.303–0.709 | 0.002** |
| **bc** | 1.282 | 1.042–1.576 | 0.051 | 2.201 | 1.408–3.440 | 0.002** |
| **bd** | 1.416 | 0.939–2.133 | 0.191 | - | - | - |
| **cc** | 0.826 | 0.503–1.354 | 0.639 | - | - | - |
| **cd** | 0.851 | 0.327–2.214 | 0.823 | - | - | - |

*Note:* a, *HTT* CAG repeat range 7-12; b, *HTT* CAG repeat range 13-20; c, *HTT* CAG repeat range 21-26; d, *HTT* CAG repeat range 27-35; OR, Odds Ratio; CI, 95% confidence interval; p, p-value; ** marking p-values < 0.01; values of this table are shown graphically in Fig. 1C; allelic combinations were taken into account if at least 5 subjects per group were available

**Supplementary Table 7. Association of disease risk with allelic combinations of *HTT* CAG repeat ranges defined by variable split points redefining ranges of b and c (≥ 48 years)**

| Split point | Repeat ranges | Genotype | MDD | | | ANX | | |
|---|---|---|---|---|---|---|---|---|
| | | | OR | CI | p | OR | CI | p |
| **17** | b: 13-16 | bb | 0.653 | 0.309–1.378 | 0.260 | 0.629 | 0.082–4.816 | 0.653 |
| | c: 17-26 | bc | 0.817 | 0.665–1.004 | 0.054 | 0.712 | 0.417–1.215 | 0.211 |
| **18** | b: 13-17 | bb | 0.860 | 0.705–1.049 | 0.137 | 0.736 | 0.440–1.231 | 0.241 |
| | c: 18-26 | bc | 1.034 | 0.868–1.231 | 0.709 | 1.036 | 0.677–1.585 | 0.872 |
| **19** | b: 13-18 | bb | 0.837 | 0.702–0.997 | 0.046* | 0.652 | 0.418–1.018 | 0.058 |
| | c: 19-26 | bc | 1.112 | 0.931–1.328 | 0.241 | 1.054 | 0.683–1.627 | 0.811 |
| **20** | b: 13-19 | bb | 0.789 | 0.663–0.939 | 0.008** | 0.538 | 0.350–0.826 | 0.004** |
| | c: 20-26 | bc | 1.276 | 1.059–1.538 | 0.010* | 1.533 | 0.991–2.373 | 0.054 |
| **21** | b: 13-20 | bb | 0.807 | 0.673–0.967 | 0.020* | 0.463 | 0.303–0.709 | < 0.001*** |
| | c: 21-26 | bc | 1.282 | 1.042–1.576 | 0.019* | 2.201 | 1.408–3.440 | < 0.001*** |
| **22** | b: 13-21 | bb | 0.827 | 0.683–1.002 | 0.052 | 0.491 | 0.319–0.756 | 0.001** |
| | c: 22-26 | bc | 1.330 | 1.058–1.671 | 0.014* | 2.352 | 1.467–3.770 | < 0.001*** |
| **23** | b: 13-22 | bb | 0.817 | 0.666–1.001 | 0.051 | 0.561 | 0.356–0.883 | 0.013* |
| | c: 23-26 | bc | 1.263 | 0.982–1.624 | 0.068 | 1.962 | 1.163–3.309 | 0.010* |

*Note:* OR, Odds Ratio; CI, 95% confidence interval; p, p-value; *, ** and *** marking nominal p-values < 0.05, < 0.01 and < 0.001 respectively; values are shown graphically in Fig. 1D

**Supplementary Table 8. Association of MDD risk with allelic combinations of *HTT* CAG repeat counts defined by variable split points**

| Split point | Repeat ranges | OR | CI | p |
|---|---|---|---|---|
| **17** | at least one allele ≥ 17 | 1.168 | 0.580 - 2.351 | 0.664 |
| | both alleles < 17 | 0.856 | 0.425 - 1.724 | 0.664 |
| **18** | at least one allele ≥ 18 | 1.132 | 0.932 - 1.376 | 0.211 |
| | both alleles < 18 | 0.883 | 0.727 - 1.073 | 0.211 |
| **19** | at least one allele ≥ 19 | 1.183 | 0.994 - 1.409 | 0.058 |
| | both alleles < 19 | 0.845 | 0.71 - 1.006 | 0.058 |
| **20** | at least one allele ≥ 20 | 1.270 | 1.067 - 1.513 | 0.007** |
| | both alleles < 20 | 0.787 | 0.661 - 0.938 | 0.007** |
| **21** | at least one allele ≥ 21 | 1.247 | 1.036 - 1.500 | 0.019* |
| | both alleles < 21 | 0.802 | 0.667 - 0.965 | 0.019* |
| **22** | at least one allele ≥ 22 | 1.241 | 1.019 - 1.512 | 0.032* |
| | both alleles < 22 | 0.806 | 0.661 - 0.982 | 0.032* |
| **23** | at least one allele ≥ 23 | 1.259 | 1.017 - 1.559 | 0.034* |
| | both alleles < 23 | 0.794 | 0.641 - 0.983 | 0.034* |

*Note:* OR, Odds Ratio; CI, 95% confidence interval; p, p-value; * and ** marking nominal p-values < 0.05 and < 0.1 respectively, results are shown graphically in Sup. Fig. 4

**Supplementary Tab. 9.1 Associations of disease status with *HTT* CAG repeats in different age groups**

| | < 48 years | | ≥ 48 years | |
|---|---|---|---|---|
| **Term** | **β** | **$P_{FDR}$** | **β** | **$P_{FDR}$** |
| **A1** | -0.030 | 0.770 | 0.046 | 0.601 |
| **A2** | -0.107 | 0.300 | 0.177 | 0.046* |
| **$A1^2$** | -0.005 | 0.961 | -0.063 | 0.472 |
| **$A2^2$** | 0.082 | 0.432 | -0.172 | 0.052 |
| **A1 × A2** | -0.292 | 0.010** | -0.010 | 0.908 |

*Note*: β, standardized beta coefficient; p, p-value; * and ** marking p-values < 0.05 and < 0.01, respectively; ×, interaction; see methods section for full model description

**Supplementary Tab. 9.2 Associations of disease status with *HTT* CAG repeats and age (entire sample)**

| Term | β | p |
|---|---|---|
| **A1** | 0.045 | 0.621 |
| **A2** | 0.183 | 0.041* |
| **$A1^2$** | -0.087 | 0.385 |
| **$A2^2$** | -0.180 | 0.052 |
| **A1 × A2** | -0.011 | 0.908 |
| **Age** | 0.366 | < 0.001*** |
| **Age × A1** | -0.050 | 0.596 |
| **Age × A2** | -0.194 | 0.037 |
| **Age × $A1^2$** | 0.060 | 0.557 |
| **Age × $A2^2$** | 0.199 | 0.040* |
| **Age × A1 × A2** | -0.189 | 0.065 |

*Note*: β, standardized beta coefficient; p, p-value; * and *** marking p-values < 0.05 and < 0.001, respectively; ×, interaction; see methods section for full model description

**Supplementary Table 10. Supplementary information about analyzed clinical variables**

| Short name | Explanation | Syntax in MARS database |
|---|---|---|
| **HMD at admission** | Hamilton Depression Rating Scale (21-item version) sum score at admission | HMD_00 |
| **BDI at admission** | Beck Depression Inventory-II sum score at admission | BDI_00 |
| **Apathy at admission** | Apathetic syndrome subscore (based on the AMDP) at admission | AMDAPA00 |
| **HAMA at admission** | Hamilton Anxiety Rating Scale sum score at admission | HMA_00 |
| **Early partial response** | Early partial response (≥25% HMD reduction after week 2) (binary variable) | HD_2WO |
| **Apathy at week 4** | Apathetic syndrome subscore (based on the AMDP) at week 4 | AMDAPA04 |
| **Apathy dif. week 0/4** | Difference in AMDP apathetic syndrome score between week 4 and admission | AMDAPA_d |
| **Remission at discharge** | Remission at discharge (HMD < 10) (binary variable) | KRM_E |
| **Age at onset** | Age at psychiatric illness onset | age_on |
| **Suicidal attempt(s)** | Suicidal attempt(s) in the medial history (binary variable) | kgsu_n |
| **Suicidality weeks 0 to 4** | Suicidal thoughts during the first 4 weeks (defined as at least one of weekly HMD item 3 measurement ≥ 2 between admission and week 4) | sui_ever_hmd3_2 |
| **Family history MDD** | Family history positive for MDD (single episode or recurrent episodes) (binary variable) | family |
| **Family history suicides** | Family history positive for suicide (binary variable) | FA_X60 |
| **Family history dementia** | Family history positive for dementia (binary variable) | FA_F00 |
| **BDI anhedonia items** | BDI-II subscore at admission including only anhedonia related items: 4; 18; 21 (4: loss of pleasure; 18: loss of appetite; 21: loss of interest in sex) | BDI0_04_18_21 |
| **BDI without anhedonia items** | BDI-II score at admission including items 1-3; 5-17; 19; 20 and excluding anhedonia related items (4; 18; 21) | BDI0_no_04_18_21 |
| **Cigarette smoking** | Cigarettes per day | Zigis |

**Supplementary Table 11. Association of *HTT* CAG repeat counts with anhedonia and addictive behavior related clinical variables in MD**

| | | | $A1_{orth}$ | | A2 | | Age | | $A1_{orth}$ × A2 | | $A1_{orth}$ × Age | | A2 × Age | |
|---|---|---|---|---|---|---|---|---|---|---|---|---|---|---|
| **Variable** | **n** | **Rsq** | **β** | **p** | **β** | **p** | **β** | **p** | **β** | **p** | **β** | **p** | **β** | **p** |
| **BDI anhedonia items** | 441 | 0.012 | -0.013 | 0.841 | -0.024 | 0.734 | 0.026 | 0.593 | 0.140 | 0.006** | -0.051 | 0.430 | 0.028 | 0.688 |
| **BDI without anhedonia items** | 405 | 0.014 | -0.015 | 0.821 | -0.012 | 0.866 | -0.064 | 0.198 | 0.125 | 0.019* | -0.062 | 0.361 | -0.002 | 0.975 |
| **Cigarette smoking** | 1025 | 0.019 | 0.034 | 0.422 | 0.027 | 0.536 | -0.127 | < 0.001*** | 0.025 | 0.456 | -0.049 | 0.238 | 0.045 | 0.288 |

*Note:* n, number of available subjects; Rsq, adjusted $R^2$; β, standardized beta coefficient; p, p-value; *, ** and *** marking nominal p-values < 0.05, <0.01 and < 0.001, respectively; BDI, Beck Depression Inventory; BDI anhedonia related items: 4, 21, 18; for further explanations of the clinical variables see Sup. Tab. 10
